# Temporal pattern recognition in retinal ganglion cells is mediated by dynamical inhibitory synapses

**DOI:** 10.1101/2023.01.12.523643

**Authors:** Simone Ebert, Thomas Buffet, B.Semihcan Sermet, Olivier Marre, Bruno Cessac

**Affiliations:** Biovision and Neuromod Institute, Université Côte d’Azur, Sophia Antipolis, France; Sorbonne Université, INSERM, CNRS, Institut de la Vision, 17 rue Moreau, F-75012 Paris, France; Institut Pasteur, Paris, France

## Abstract

A fundamental task for the brain is to generate predictions of future sensory inputs, and signal errors in these predictions. Many neurons have been shown to signal omitted stimuli during periodic stimulation, even in the retina. However, the mechanisms of this error signaling are unclear. Here we show that depressing inhibitory synapses enable the retina to signal an omitted stimulus in a flash sequence. While ganglion cells, the retinal output, responded to an omitted flash with a constant latency over many frequencies of the flash sequence, we found that this was not the case once inhibition was blocked. We built a simple circuit model and showed that depressing inhibitory synapses were a necessary component to reproduce our experimental findings. We also generated new predictions with this model, that we confirmed experimentally. Depressing inhibitory synapses could thus be a key component to generate the predictive responses observed in many brain areas.

## 1 Introduction

A long standing hypothesis is that visual neurons do not signal the visual scene *per se*, but rather surprising events, eg. mismatches between observation and expectation formed by previous inputs [3]. It has been observed in a number of sensory modalities that neurons strongly respond when a sequence of repetitive stimuli is unexpectedly interrupted [35, 4, 21]. In the retina, this phenomenon has been coined the Omitted Stimulus Response (OSR) [30]. When a periodic sequence of flashes suddenly ends, some ganglion cells emit a large response. Interestingly, the latency of this response shifts with the period of the flash sequence, so that the ganglion cell responds to the omitted flash with a constant latency. This suggests that the retina forms predictions of observed patterns, and responds to a violation of its internal expectation. However, the mechanisms by which the retina achieves this remain unclear and debated [29, 39, 9, 32].

Here we investigated how inhibitory amacrine cells affect the OSR and showed that depression in inhibitory synapses can account for this characteristic latency shift. To this end, we performed electrophysiological recordings of retinal ganglion cells and found that blocking inhibitory transmission from glycinergic amacrine cells selectively abolished the predictive latency shift of the OSR. To better understand how glycinergic inhibition impacts the latency of the OSR, we developed a circuit model equipped with a glycinergic amacrine cell. This model could reproduce the latency shift of the OSR when the glycinergic synapse showed short-term depression, thereby adjusting its weight to the stimulus frequency. Our model generated several predictions about the OSR, which we could confirm in experiments. The latency shift that is characteristic of the OSR is thus due to a depressing inhibitory synapse whose weight is changed by the stimulus frequency. For low frequency sequence, the synaptic weight is large and this increases the latency of the response, while for high frequency stimuli, the weight is low due to depression, and the latency is only shifted by a small amount. Our results suggest a generic circuit to generate responses to surprise that could be implemented in several brain areas.

## 2 Results

### 2.1 ON biphasic ganglion cells exhibit an Omitted Stimulus Response to dark flashes

Using a multi-electrode array of 252 electrodes, we extracellularly recorded the spiking activity of ganglion cells from the mouse retina. [20] (Figure 1 A). We presented sequences of 12 full-field dark flashes of 40 ms duration each, at frequencies of 6, 8, 10, 12 and 16 Hz, with a grey baseline illumination. We estimated the receptive fields of the same retinal ganglion cells with a checkerboard-like noise. We defined ON cells based on their receptive fields (responding to a light increase, 1 B). We focused on the response of ON cells to sequences of dark flashes.

**Figure 1:**
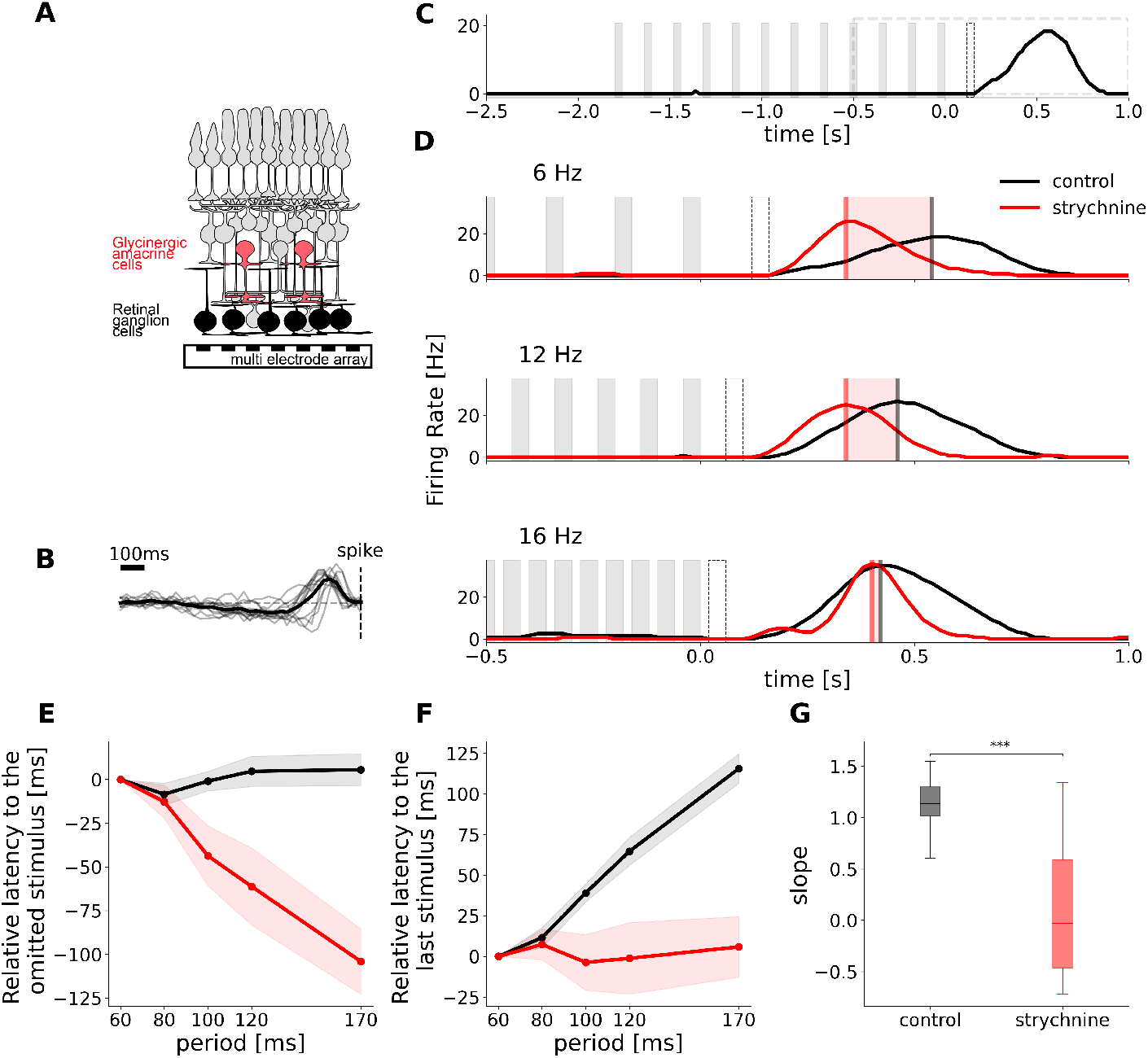
Glycinergic Amacrine cells are necessary for predictive timing of the OSR. **A**. Schematic representation of the retina, with the activity of retinal ganglion cells being recorded with a multi electrode array. **B**. Temporal traces of the receptive fields of the cells that exhibit an OSR. **C**. Example of OSR. The cell responds to the end on the stimulation, after the last flash. The times of dark flashes are represented by grey shaded rectangles. The black dotted rectangle shows the timing of the omitted flash. The grey dotted rectangle shows the time period of focus in panel D. **D**. Experimental recording of the OSR in one cell in control condition (black) and with strychnine to block glycinergic amacrine cell transmission (red). Firing rate responses to flash trains of 3 different frequencies are aligned to the last flash of each sequence. Flashes are represented by gray patches, vertical lines indicate the maximum of the response peak, red shaded areas indicate the temporal discrepancy between control and strychnine conditions. **E**. Mean latency between OSR and the omitted flash plotted against the period of the stimulus for a population of *n* = 12 cells. Latency is expressed relatively to the latency of the response to the 16Hz stimulus in the control condition. **F**. Mean latency between OSR and last flash in the stimulus plotted against stimulus period. Control latencies shift with the period of the stimulus wit a slope 1.143 *±*0.08. With strychnine this shift is abolished (slope = 0.07*±* 0.18). Latency is expressed relatively to the latency of the response to the 16Hz stimulus in the control condition. **G**. The slope of the latency shift decreases significantly when strychnine is added, p-value = 0.0004 (paired t-test).

Many ON cells responded with a broad peak of activity after the stimulus stopped for all frequencies tested (*n* = 74). A subset of these cells (36 %, n = 26, fig. 1 D, see Methods) shifted the response latency according to the stimulus frequency. They exhibit an “Omitted Stimulus Response” (OSR), as it was first described in [30]: when the period of the flash train increases, the latency of the response to the last flash in the stimulus shifts by the same amount (fig. 1C,D), such that the latency of the response to the omitted stimulus is constant (fig. 1E). This indicates that the retina has a precise temporal expectation of when the next flash should have occurred and shifts the latency of its response accordingly. This is illustrated in fig. 1F, where the relation between the latency of the response and the period of the flash sequence is linear, with a slope of nearly 1 (fig. 1G). In the following we will refer to this specific relation as “latency shift”. Interestingly, the ON cells showing this latency shift mostly had a biphasic response profile 1 B).

### 2.2 Amacrine cells are required for the latency shift in the Omitted Stimulus Response

It remains unclear how the retinal circuit generates the OSR. Previous studies have shown that the ON bipolar cell pathway is necessary to have a response *per se* [30], but the components of the retinal circuit needed for the latency shift are yet to be determined. We hypothesized that the inhibitory cells of the retinal circuit are responsible for the shift in latency, as inhibition has been shown to shift latency in various circuits [34, 37, 38, 8]. Amacrine cells are the main class of inhibitory interneurons in the mouse retina. To investigate this further, we blocked glycinergic transmission using strychnine (2*µM*) and recorded the spike responses of retinal ganglion cells to flash trains of varying frequencies (see Methods). As glycinergic transmission is only employed by certain classes of inhibitory amacrine cells in the mouse retina [40] this blocks only a subset of amacrine cells.

While the response after the sequence end remained after strychnine application, we observed that the slope between response latency and stimulus period decreased from an average of 1.13 *±*0.08 in the control condition to 0.07 *±*0.18 after strychnine was added (Figure 1 F and G, *n* = 12, see Methods). While the OSR occurred at roughly the same time in the highest frequency tested, the peak was significantly advanced after low frequency flashes compared to the control condition (Figure 1 D). As a consequence, the OSR did not have a constant latency relative to the omitted stimulus after strychnine was added (Figure 1 E). These results demonstrate that glycinergic amacrine cells are a key contributor to the OSR. Although they do not generate the response alone, they are crucial for the latency shift, which is a hallmark of the OSR.

### 2.3 A circuit model with depressing synapses in inhibitory glycinergic amacrine cells explains the latency shift

Our experimental results provided compelling evidence that glycinergic amacrine cells provide inputs which are required to achieve the latency shift of the OSR. However, it remained unclear how the retinal circuit exhibits an OSR in a glycinergic dependent way. We developed a mechanistic model in which we explicitly simulated inputs from glycinergic amacrine cells to understand their role in the latency shift of the OSR.

Since we focused on biphasic ON ganglion cells, we equipped our model with two ON inputs, one being excitatory *E*^*ON*^ and mimicking ON bipolar cell input, and one being inhibitory *I*^*ON*^, conveying broad delayed inhibition. This delayed inhibitory input summarizes the influence of various inhibitory pathways (horizontal cells, GABAergic amacrine cells, ON glycinergic amacrine cells) that generate the biphasic response profile (see discussion). In addition, we explicitly included a glycinergic amacrine cell with an OFF polarity, 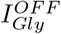, in order to provide inhibition to dark stimuli. All three units receive the visual stimulus as input, and connect onto a ganglion cell *G* (Fig. 2 A, see Methods for details).A characteristic feature in the Omitted Stimulus Response is that the latency shifts by the same amount as the stimulus period. It has been shown that the relative strengths of excitation and inhibition can determine the latency of the response [34, 37, 38, 8]. We reasoned that the effective strength of the inhibitory input should thus depend on the frequency of the flash sequence. This can be achieved with a dynamic synapse, i.e. a synapse whose strength varies with the frequency of the stimulation.

**Figure 2:**
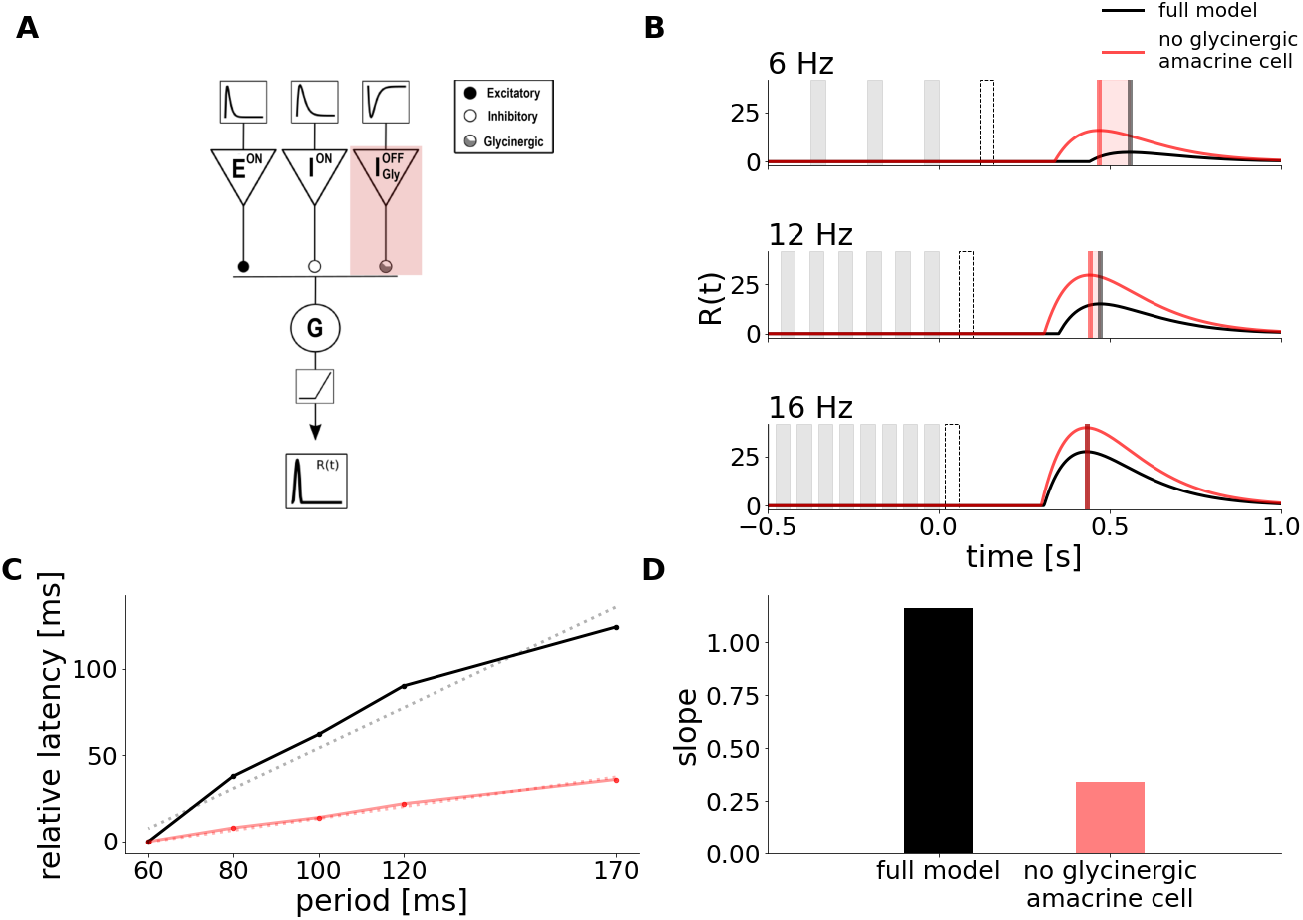
Mechanistic model replicates latency shift and strychnine experiment. **A**. Schematic description of the Model. It is composed of an ON excitatory input *E*^*ON*^, an ON inhibitory input *I*^*ON*^, and an OFF inhibitory input *I*^*OF F*^ representing a glycinergic amacrine cell. Each of those units receives as input the convolution of the stimulus with a monophasic temporal kernel, determining the cells polarity, and connects onto a ganglion cell *G*. The response of *G* is then passed through a nonlinearity to simulate the cells’ firing rate. The synapse from 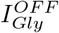 to *G* can adapt its strength to the stimulus via short-term depression. The shaded red area represents the weight of this glycinergic amacrine cell being set to zero to simulate the effect of strychnine. **B**. Simulation of the model responses to flash trains of 3 different frequencies with the full model (black, control), and the weight of 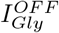 set to 0 (red, strychine). The weight of *I*^*ON*^was decreased in this simulation, accounting for the broad effect of strychnine, which likely reduces inhibition overall. The times of dark flashes are represented by grey shaded rectangles, while the black dotted rectangle shows the timing of the omitted flash. **C**. Latency of the OSR plotted against stimulus period in control and strychnine simulation. The latency is expressed relatively to the latency of the response of the full model to the 16Hz stimulus. The slope of the latency shift decreased from 1.16 to 0.34 when 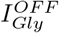 was set to 0 (dotted lines).**D**. Value of the slope fitted to latency shift in the full model and strychnine simulation.

Several previous reports have shown that inhibitory synapses can be depressing, i.e. have a decreasing weight depending on previous inputs [18, 36, 23, 17, 14]. It has been hypothesized that the variation in synaptic strength results from varying availability of vesicles in the readily releasable vesicle pool, which gets gradually depleted upon persistent inputs [31, 5, 25]. In the retina, this has been modelled via dynamical systems of vesicle pools, where the value of one of the variables directly serves as input to the postsynaptic cell [26, 28]. To keep our model as simple as possible, we modelled the glycinergic synapse as a depressing synapse using one kinetic equation to simulate synaptic vesicle occupancy, similar to cortical models of short-term plasticity [33, 42, 11], but replaced spiking inputs by continous presynaptic voltage. The vesicle occupancy then scales the output of the cell (see Methods).

This mechanistic model allowed us to reproduce the main properties of the recorded ganglion cells. Thanks to excitatory and inhibitory ON inputs, it has an ON biphasic impulse response. It also responds with a peak at the end of dark periodic flash stimuli, due to delayed disinhibition, because inhibition has a slower temporal filter than excitation (Figure 2 B). Our model also successfully simulated an OSR whose latency with respect to the last flash increased with the stimulus period with a slope around 1, and thus with a constant latency with respect to the omitted stimulus (Figure 2 B and C, black line).

We next simulated the experimental effect of strychnine with the model by removing the glycinergic amacrine cell input. Since the ON inhibitory cell of our model also includes the effect of ON glycinergic amacrine cells, we also decreased the weight of the ON inhibitory input (note however that our results did not depend on that additional modification, see discussion). Figure 2 B and C (red) show that the model replicates the experimental effect of strychnine. The slope of the latency shift decreases from 1.16 to 0.34 when 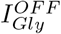 is removed from thecircuit.

To test whether the model achieves the latency shift of the response peak thanks to the dynamical synapse, we simulated the response of the model while keeping the occupancy of the glycinergic amacrine cell synapse constant at 1. (3 A). The latency increased in all frequencies simulated, but the slope of the latency shift decreased to 0.32 (see 3 B-D). In our model, the dynamical synapse is thus essential to achieve the latency shift of the response with a slope of 1 observed experimentally. Note that there was also an overall increase in latency which can be explained by the fact that the fixed synapse is stronger than the dynamic counterpart, since it is never at full occupancy when stimulated due to depression.

### 2.4 The depressing inhibitory synapse induces a latency shift

To understand how the model can account for the latency shift in the OSR, it is helpful to look at the temporal evolution of its internal variables. The ON excitatory input evokes a hyperpolarization in response to dark flashes, and cancels the depolarization evoked by the slightly slower ON inhibitory cells during the flash sequence. At the end of the flash sequence, due to differing time constants, there is a time window where depolarization due to the inhibitory delayed cell exceeds the hyperpolarization due to the bipolar cell, and triggers a spiking response (Figure 4 A). This is similar to many classical rebound responses recorded experimentally, and it can be predicted with a biphasic filter. But by itself, this biphasic filter would not predict the latency shift as observed experimentally. As we will describe in the following paragraph, this latency shift is due to the specific effect of the glycinergic amacrine cell equipped with a depressing synapse.

**Figure 3:**
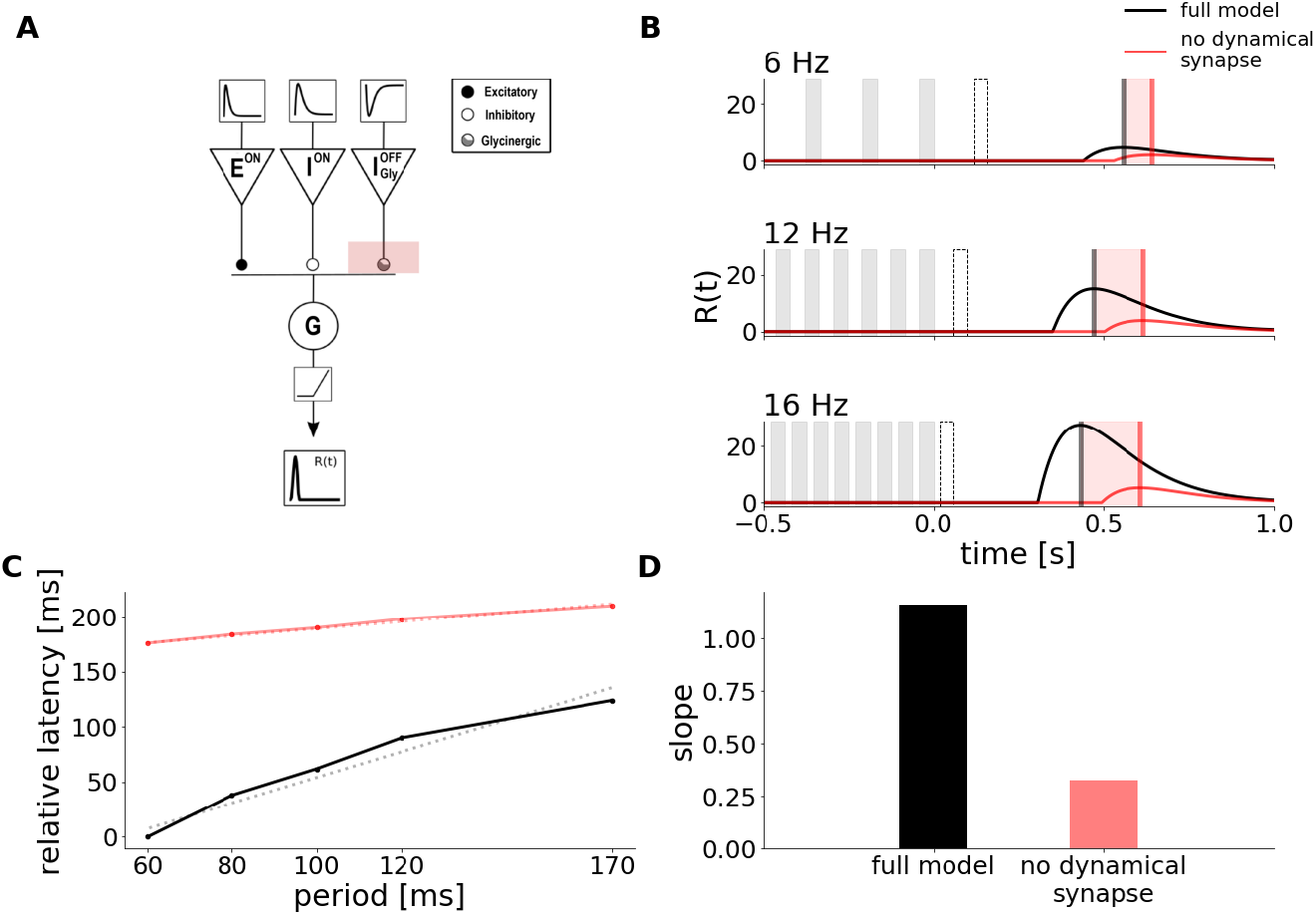
Short-term plasticity is the crucial component for latency shift **A**. Schematic description of the Model, as in Figure 2. The shaded red area now represents removing the dynamic characteristic of the glycinergic synapse. **B**. Simulation of the Model responses to flash trains of 3 different frequencies with dynamic occupancy (black) and the weight of 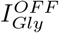 held constant (red). The times of dark flashes are represented by grey shaded rectangles, while the black dotted rectangle shows the timing of the omitted flash. **C**. Latency of the OSR plotted against stimulus period in control and strychnine simulation. The latency is expressed relatively to the latency of the response to the 16Hz stimulus in the control condition. The slope of the latency shift decreased from 1.16 to 0.32 when the weight of 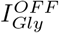 was held constant (dotted lines). **D**. Value of the slope fitted to latency shift in the full model and without the adaptive property of the glycinergic synapse.

**Figure 4:**
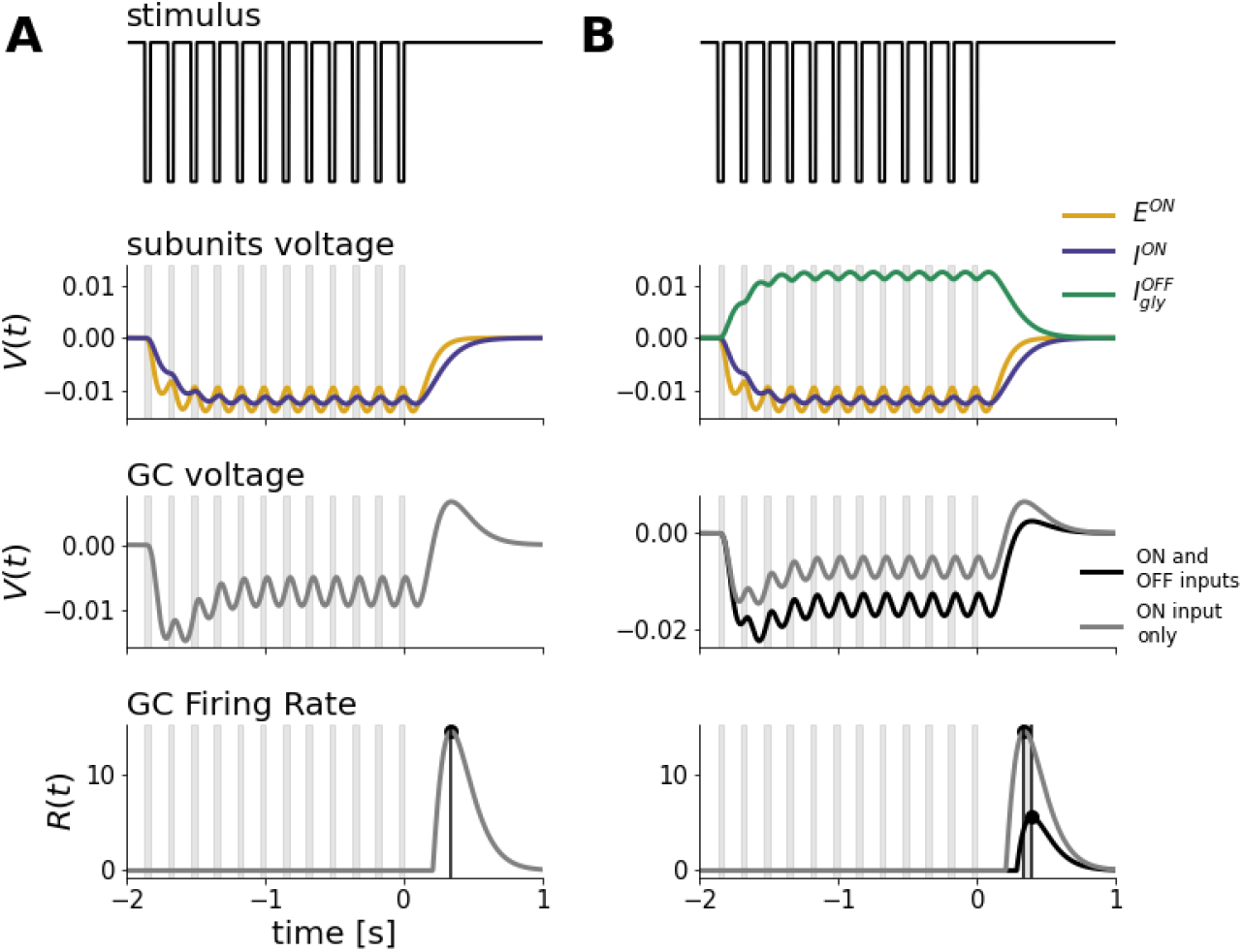
ON components of the model produce a peak after stimulus end while the glycinergic OFF input shifts the latency. **A. - B**. Internal model responses to a 6 Hz dark flash stimulus. The glycinergic OFF synapse is dynamic. From top to bottom: Stimulus Intensity, Bipolar and Amacrine voltage responses, ganglion cell voltage and firing rate. **A**. ON excitation and inhibition hyperpolarize in response to dark flashes. When both inputs are substracted in the Ganglion cell, its voltage sum hyperpolarizes during flash presentation, followed by an overshoot of disinhibition due to the slower response profile of the inhibitory input. After passing the voltage through a rectification function, only the disinhibitory peak after stimulus end remains in the firing rate. **B**. The voltage of the glycinergic OFF input depolarizes in response to dark flashes, passing additional inhibition onto the ganglion cell. This lowers the GC voltage response and increases the latency between peak and stimulus end in the firing rate. Last two panels compare the models’ simulation and peak time-point with (black) and without (grey) the input from 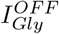.

Since this glycinergic amacrine cell is an OFF cell, it inhibits the ganglion cell in response to dark flashes. This has the effect of delaying the spiking response at the end of the flash sequence and to increase its latency (Figure 4 B, black compared to grey). The latency of the response is then shifted for different stimulus frequencies because the depressing synapse changes the strength of the glycinergic inhibition. In this synapse, the vesicle occupancy represents the amount of synaptic depression and decreases when stimulation starts (Figure 5 A, 3rd row). It then reduces the current input from 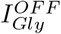 to the ganglion cell (Figure 5 A, 4th row). This reduction of inhibition shifts the OSR towards an earlier response, reducing its latency (Figure 5 A, 5th row).

**Figure 5:**
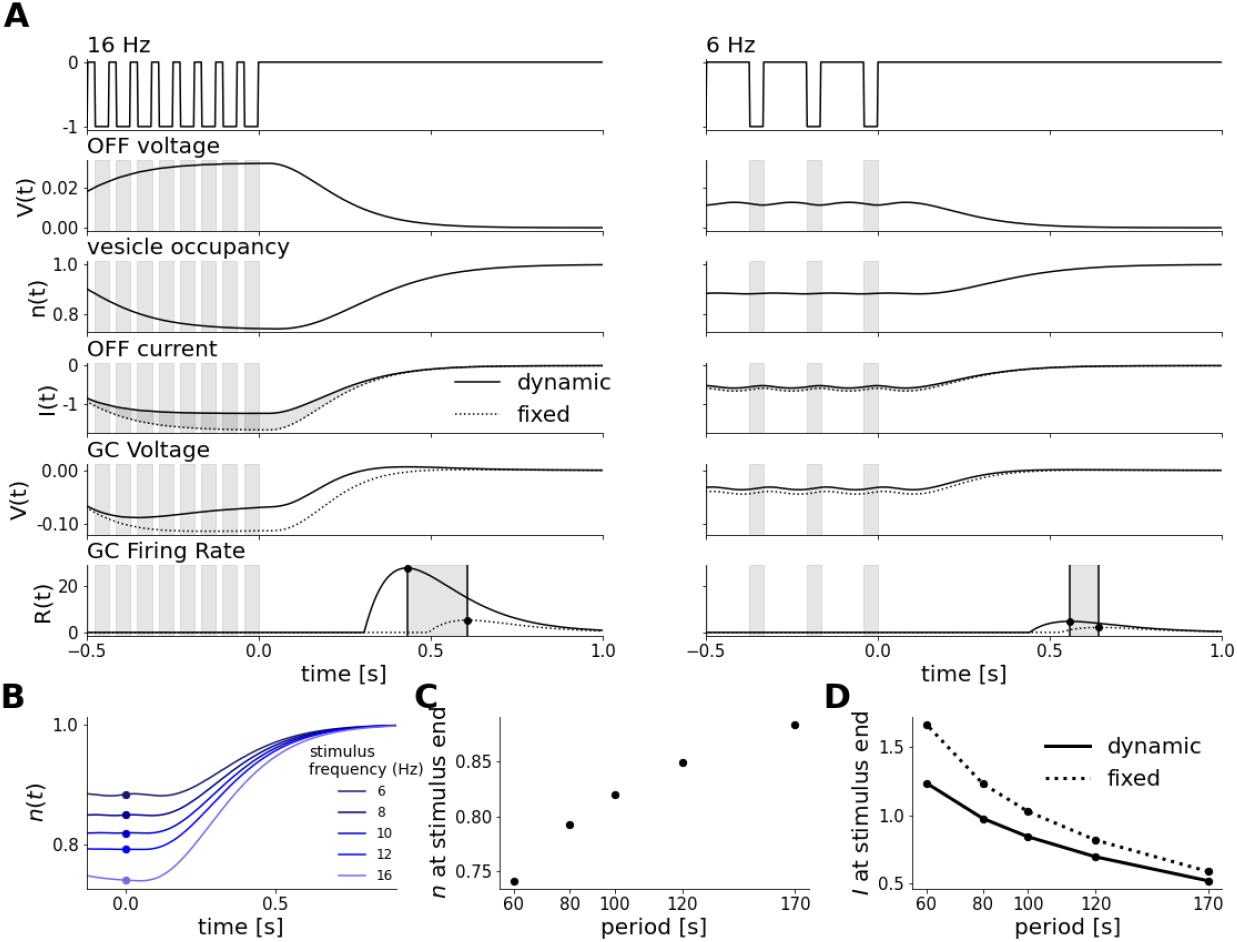
Synaptic depression scales OFF glycinergic input to stimulus frequency and thereby shifts the latency of the response. **A**. Impact of occupancy scaling on 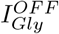 current input to *G* for a fast (16 Hz, left) and a slow (6 Hz, right) stimulus. From top to bottom: Stimulus Intensity, 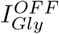 voltage, vescile occupancy, current input, *G* voltage and firing rate. Last 3 panels compare simulations with dynamic occupancy (solid lines) to when the occupancy is held constant (dotted lines). Depression has the effect to advance the OSR peak, more so for fast than slow frequencies. **B**. Occupancy traces to flash stimuli of different frequencies aligned to the last flash. Dots indicate the occupancy level at stimulus end. **C**. Level of occupancy after stimulus end scales with the period of the stimulus. **D**. 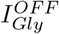 current input is decreased by short-term depression, more so for fast than slow frequencies. Dotted line shows current with fixed occupancy, solid line with dynamic occupancy.

Fast frequency stimuli cause stronger depression, reducing the 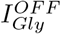 current input by 30 %. This has a strong impact on the latency, which is more than 100 ms shorter when the synapse is depressed. In contrast, slow frequency stimuli cause only weak depression, reducing the *A*_*Gly*_ by only10 %. The latency was thus only slightly reduced in that case. (compare 5 A and B).In summary, the steady state vesicle occupancy of the synapse is determined by the stimulus frequency (5 C and D). The vesicle occupancy can then reduce the inhibitory current, yielding a large reduction for fast inputs and a small reduction for slow inputs (5 E). As a result, vesicle occupancy acts like a scaling factor to tune the inhibitory current input and thereby shifts the response latency based on stimulus frequency. This explains how the latency shift observed experimentally is achieved via glycinergic inputs.

### 2.5 The depressing inhibitory synapse predicts other features of the Omitted Stimulus Response

Can our model give other predictions about the OSR ? Since the glycinergic inhibitory synapse is more depressed at high frequency, the OSR is less inhibited, and its amplitude is thus stronger compared to low frequencies (compare Figure 5 A and B, 5th row). This trend was also observed in our experiments. Both simulated and experimental amplitudes showed a negative correlation between the response amplitude and the stimulus period (*−* 0.87*±* 0.02, *n* = 14) in data and as well *−* 0.87 in simulations).

An important consequence of our depressing synapse model is that it takes several flashes to reach steady state in the vesicle occupancy. If we shorten the flash sequence, the vesicle occupancy will not reach that steady state, and this should have predictable consequences on response amplitude and latency. We simulated the response to long flash trains consisting of 12 flashes (as in the experiments and simulations above) and shorter sequences of only 5 flashes.

Our simulations predicted that the amplitude of the OSR decreases when the stimulus contains only 5 flashes. We tested that in experiments and found the same tendency: the OSR amplitude was significantly smaller in all but the lowest frequency tested (see Figure 6 A)

**Figure 6:**
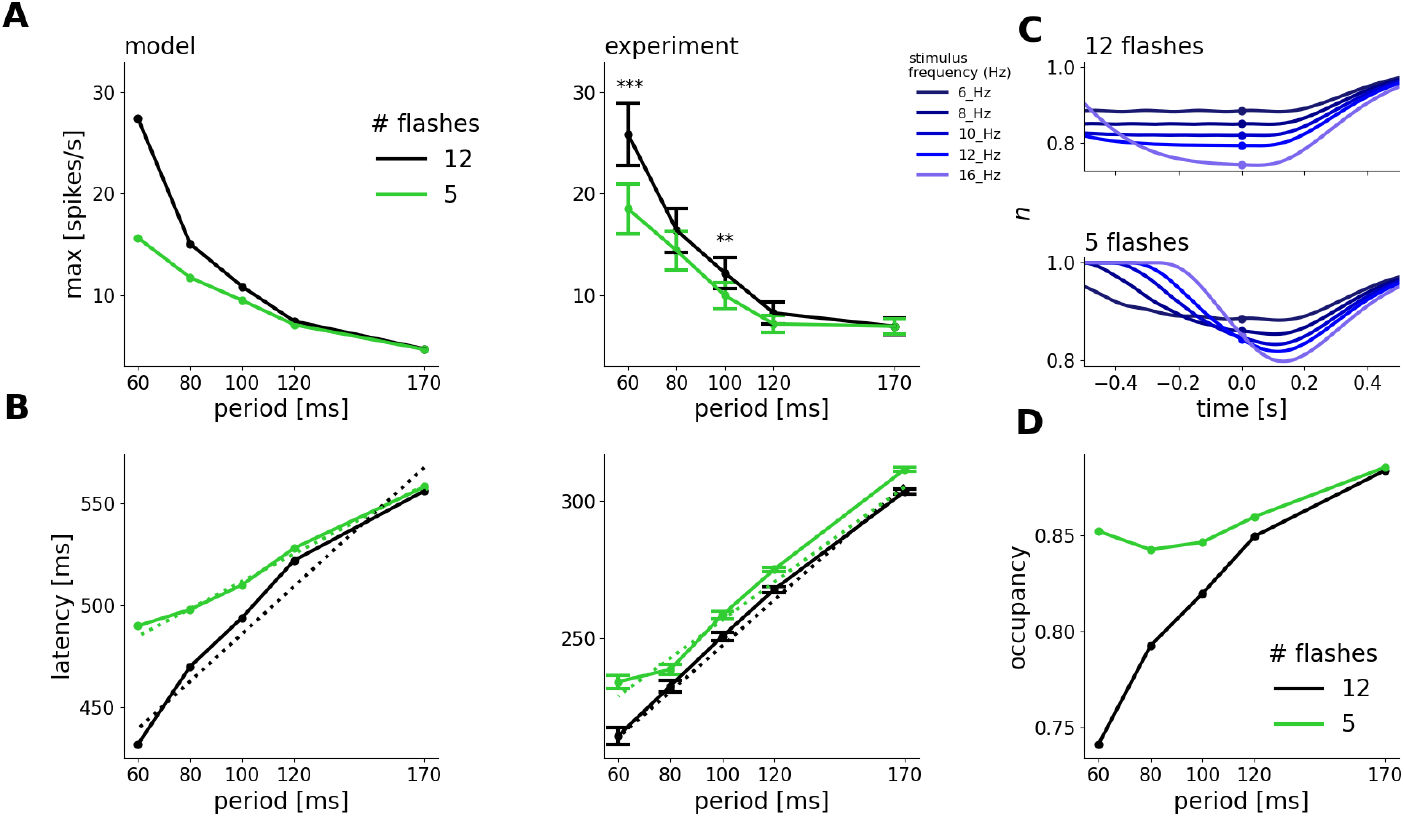
Latency shift decreases for shorter stimuli because of lacking steady state occupancy. **A**. Amplitude of the OSR against stimulus period for 12 and 5 flashes in the stimulus in simulations (left) and experiments (right). Amplitudes to 6 Hz and 10 Hz stimuli were significantly different after Bonferroni-Holm correction [12] (6 Hz: *p* = 0.000006, 10 Hz: *p* = 0.001). **B**. Latency against stimulus period in simulations (left) and experiments (right). Simulated slopes decreased from 1.16 after 12 flashes to 0.67 after 5 flashes. Experimental slopes decreased from 0.84*±* 0.02 to 0.69 *±*0.04. The latency after 12 and 5 flashes was significantly different for 10 Hz (*p* = 0.03), 12 Hz (*p* = 0.04) and 16 Hz stimuli (*p* = 0.02). **C**. Temporal traces of vesicle occupancy to all frequencies simulated, for 12 Flashes (upper) and 5 Flashes (lower). Dots indicate occupancy at stimulus end. Traces do not reach a steady state for 5 flashes. **D**. Scaling of occupancy with stimulus period in 5- and 12-flash scenario.

Another model prediction was that the slope of the relation between OSR latency and stimulus period should decrease for shorter flash trains, reaching only a value of 0.67 for 5 flashes, compared to 1.16 when 12 flashes were presented (see Figure 6 B, left) In our model, this is a consequence of the dynamics of the depressing synapse.

In the 5-flash scenario, our model predicted that the stimulus is too short for the synapse to reach a steady state occupancy when the stimulus frequency is high (Figure 6 C). 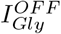 hence provides a larger inhibitory input than for a longer sequence. This increases the response latency and changes the slope of the relation between latency shift and stimulus period (Figure 6 D).

In our experiments, while there was no difference for low frequency stimuli, the absolute latency of the OSR was much larger after 5 flashes than after 12 when the stimulus frequency was high (see Figure 6 B). This change in latency led to a reduction of the slope value from 0.84 *±*0.02 to 0.69*±* 0.04 in experiments, consistent with the model prediction. The agreements between the model predictions and experiments provide further evidence for the validity of our model, and for the key role of a depressing inhibitory synapse.

## 3 Discussion

The Omitted Stimulus Response is an example of sophisticated feature detection that takes place already in the retina. This phenomenon implies that retinal ganglion cells can carry a dynamic prediction of their future visual input with high temporal precision, and selectively respond when this prediction is not matched.

With this work, we provide evidence that the latency shift of the OSR, which allows a constant latency relative to the omitted stimulus, is generated by inhibition from glycinergic amacrine cells. Using computational modelling, we show how inhibition can enable retinal ganglion cells to respond to the missing flash at the end of a sequence. Short-term depression in inhibitory synapses allows shifting the latency of this response.

Previous experimental studies [30, 29, 39] reported that the OSR is found in a higher proportion of retinal ganglion cells than what we observed in this study. This difference could come from the fact that we define OSR as a response with a latency shift having a slope of at least 0.7, while it is not clear whether previous studies took multiple frequencies into account when classifying the OSR. Schwartz et al. [29] also showed that blocking inhibition from amacrine cells had no effect on the OSR. But again, this study only investigated the presence or absence of the OSR under amacrine blockade, and did not investigate if the latency of the OSR shifted with the stimulus frequency. In addition, some previous studies were mostly carried out in salamander, where the underlying mechanisms may be different from the mouse.

Several theoretical models have been proposed to elucidate the mechanisms behind the Omitted Stimulus Response. Werner and Passaglia [39] proposed a dual LN-model with biphasic ON-OFF pathway interactions, which accurately captures the response peak after stimulus end via the rebound phase of the pathway selective to the opposite polarity than the stimulus. However, it fails to shift the peak latency as a function of the stimulus period with a slope of 1, which is a defining feature of the OSR. This slope value is necessary to have a response of constant latency with respect to the omitted stimulus.

A slope value of 1 indicates that the cell responds with a constant latency to the omitted stimulus, while a slope value of 0 indicates a response with a constant latency relative to the last flash. This slope value is thus a defining feature of the OSR.

When removing the depressing synapse, our model is amenable to a biphasic LN model, since the responses of our intermediate units are then linear and could be represented by a single linear filter as well. Our model then simulates the OSR in the same manner as this previous study, but the depressing inhibitory synapse was necessary to obtain the slope of 1, which is the signature of a predictive latency shift.

Gao and Berry [9] proposed intrinsic oscillatory activity in ON bipolar cells that evoked a latency shift via resonance tuned to the stimulus frequency. However, such oscillatory activity was not found in bipolar cells [7]. Our experiments where we blocked glycinergic transmission, while leaving bipolar cells intact, show that intrinsic properties of bipolar cells are not sufficient to generate a predictive latency shift.

More recently, Tanaka et al. [32] proposed that the OSR with its latency shift can arise in a deep neural network model via summation of multiple excitatory inputs with different time constants. It is difficult to evaluate whether the model accurately captures the latency shift as observed in experiments, with the correct slope value. The explanation behind this model is that the OSR latency is determined by the sum of 2 ON bipolar cells which are activated only by certain stimulus frequencies due to different temporal filtering. This purely excitatory mechanism of latency scaling is not in line with our experimental findings, suggesting that amacrine cells likely contribute to temporal filtering as well. Our hypothesis is thus that the components they isolated correspond to a mix of bipolar and amacrine cell properties.

In contrast to those previous models of the OSR, we explicitly included an inhibitory input whose contribution to the peak latency is dependent on the stimulus frequency via short-term plasticity. By doing so, we can propose a mechanistic explanation and match the latency shift of the OSR as well as various other response properties of the experimentally observed OSR.

In order to realistically simulate the models response with glycinergic amacrine cells blocked, we had to decrease the weight of the inhibitory ON input *I*_*ON*_ in our simulations. Leaving the weight of this input untouched while setting 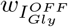 to 0, we still obtain a decrease in latency shift but this configuration generated a response to each flash of the sequence, something we did not observe experimentally. We therefore deemed this configuration as less realistic than decreasing also the weight of the ON inhibitory cell, since strychnine is likely to affect glycinergic ON inhibition as well. The components of our circuit might represent several cell types pooled together, and more detailed circuit models might give similar predictions. For example, we chose to simulate synaptic depression via a modified version of cortical STP-models with only 2 parameter rather than the more complex systems used in the retina previously [28]. We aimed at finding the minimal components necessary to obtain our results.

Dynamical synapses have previously been proposed to enable neuronal circuits in the retina to form expectations of future inputs [13] and are thus a plausible candidate to play an important role in the OSR. Previous works have shown that inhibitory synapses can be depressing [16, 15, 23]. In particular, glycinergic synapses that input to bipolar cells can be depressing [14]. However, there is no method to experimentally remove the depressing nature of the synapse without affecting the inhibitory weight, so we could not show experimentally that the depressing nature of the synapse is necessary to the OSR. However, our model predicted that the depressing inhibitory synapse should have several functional consequences, that we verified in the data. In particular, a key prediction of the depressing synapse is that the OSR requires a long enough flash sequence to accurately shift the latency, which coincides with the time needed to reach a steady state in the synaptic weights, and we confirmed this prediction experimentally.

Ultimately, our results might be of relevance to understand neuronal mechanisms of predictive coding beyond the retina. Very similar surprise responses exist in other sensory domains, such as the mismatch negativity response in the auditory cortex [24, 10, 35, 19], where neural activity is enhanced following a ‘deviant’ tone in a sequence of ‘standard’ tones. A recent study suggested that synaptic adaptation could be a key contributor to this phenomenon [2]. Following the predictive coding theory, one possible explanation is that this response emerges from an interaction between feed-forward and feedback connectivity [27, 22]Here we show that a purely feed-forward micro-circuit can generate this response to a violation of prediction via an interplay of excitation and inhibition, where synaptic depression takes place in inhibitory connections. All the components used in this micro-circuit are generic and can be found in other sensory areas [6], [33],[1], and it is thus likely that a similar circuit could be at work at the cortical level, for more complex pattern recognition than full-field flashes.

## 4 Methods and Materials

### 4.1 Experimental Setup

#### 4.1.1 Recordings

Recordings were performed on C57BL6/J adult mice of either sex. Animals were killed according to institutional animal care standards. The retina was isolated from the eye under dim illumination and transferred as quickly as possible into oxygenated Ames’ medium (Merck, A1420). The retina was extracted from the eye cup and lowered with the ganglion cell side against a multi-electrode array whose electrodes were spaced by 30*µ*m, as previously described. [20] During the recordings, the Ames’ medium temperature was maintained at 37°C. Raw voltage traces were digitized and stored for off-line analysis using a 252-channel preamplifier (MultiChannel Systems, Germany) at a sampling frequency of 20kHz. The activity of single neurons was obtained using Spyking Circus, a custom spike sorting software developed specifically for these arrays. [41]

#### 4.1.2 Visual stimulation

Visual stimuli were presented using a white LED and a Digital Mirror Device (DMD). Flash sequences contained 5 or 12 flashes of 5 different frequencies (6Hz, 8Hz,10Hz,12Hz,16Hz). Polarities were either switched from grey to black (dark flashes) or from grey to white (bright flashes). 60 trials were conducted for each stimulus, with 2-4s between each trial. The order of magnitude of the background illumination was 10^6^ R*.

#### 4.1.3 Pharmacology

To block glycinergic transmission, we dissolved strychnine (Sigma-Aldrich, S8753) in Ames’ medium at a concentration of 2*µM*, and perfused the retina with the solution at least 15 minutes before the recording.

#### 4.1.4 Latency Analysis

To determine slope of latency shift, we measured the latency between the peak firing rate and the the end of the last flash in the stimulus for all frequencies tested. We plotted these latencies against the respective period of the stimulus and fitted a straight line to determine the slope of the latency shift. Cells where classified as having an OSR in the control condition when the slope what at least 0.7 or higher. All cells where the peak time point could not be unambiguously determined in any condition were excluded from the analysis.

### 4.2 Modeling

The 3 pathways of Fig. 2 receive an input from the Outer Plexiform Layer (OPL) written as a temporal convolution of the OSR stimulus, *s*(*t*) with a linear filter of the form:

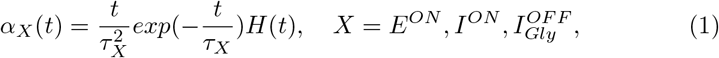

where *τ*_*X*_ is cell X characteristic time of integration (in *s*) and *H*(*t*) the Heaviside function.Thus, the inputs read:

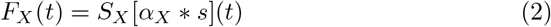

where *S*_*X*_ is a scale factor and*∗* the space-time convolution. If *S*_*X*_ is negative, *X* is an OFF cell. Note that, the stimulus being spatially uniform, the space integration reduces to a constant, so that the detailed shape of the spatial RF plays a trivial role. The OPL response is then integrated into all pathways via a linear dynamical system:

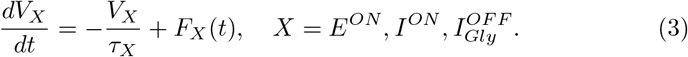

where *V*_*X*_ is the voltage of cell *X* (in Volt).

Next, all pathways provide input to the ganglion cell *G*:

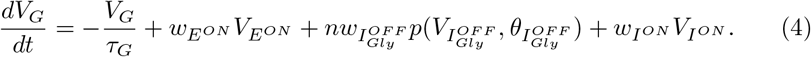

where 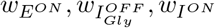 are synaptic weights (in Hz). Voltages are rectified before integrated in the ganglion cell membrane potential via :

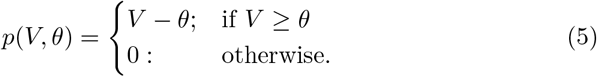

where *θ* is a threshold (in Volts).

The synaptic weight from 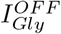 to *G* is modulated by a dimensionless variable *n*, used to simulate synaptic short-term plasticity. *n*, which interprets as a vesicle occupancy in the glycinergic amacrine synapse, obeys the kinetic equation [11] :

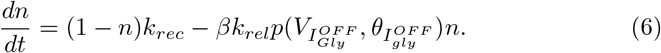

*k*_*rec*_ and *k*_*rel*_ are rate constants (*Hz*) for vesicle release and replenishment and *β* (*V* ^*−* 1^) is a scaling factor. Finally, the voltage response is passed through the piece-wise linear function *p* to obtain the firing rate.

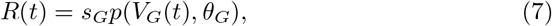

where *s*_*G*_ is a scaling factor.

Parameter values where then chosen such that simulations match the mean latencies and amplitudes of the OSR response observed in experiments and are listed in 1.

**Table 1:** Model parameter values used in simulations

| Parameter | Value | Unit |
| --- | --- | --- |
| $\tau_{EON}$ | 0.05 | $s$ |
| $\tau_{ION}$ | 0.08 | $s$ |
| $\tau_{I_{Gly}^{OFF}}$ | 0.08 | $s$ |
| $\tau_G$ | 0.1 | $s$ |
| $w_{EON}$ | 50.0 | $Hz$ |
| $w_{ION}$ | -95.0 | $Hz$ |
| $w_{I_{Gly}^{OFF}}$ | -82.0 | $Hz$ |
| $S_{EON}$ | 1.0 | $Vs^{-1}$ |
| $S_{I_{Gly}^{OFF}}$ | -0.625 | $Vs^{-1}$ |
| $S_{ION}$ | 0.625 | $Vs^{-1}$ |
| $\theta_{I_{Gly}^{OFF}}$ | 0.0 | $V$ |
| $k_{rel}$ | 4.5 | $Hz$ |
| $k_{rec}$ | 1.0 | $Hz$ |
| $\beta_{EON}$ | 13.6 | $V^{-1}$ |
| $\theta_G$ | 0.0 | $V$ |
| $s_G$ | 2200 | $HzV^{-1}$ |

## 5 Acknowledgments

We thank Romain Brette, Matthias Henning and Romain Veltz for helpful discussions about the model, and Matias Goldin and Brice Bathellier for critical reading of the manuscript. This work has been funded by a PhD fellowship from the Neuromod Institute, Université Côte d’Azur to S.E., by an ERC grant (No 101045253, DEEPRETINA) to O.M., ANR grants (DECORE, ANR-18-CE37-0011, and PerBaCo, ANR-22-CE37-0016-02) to O.M., a grant from Retina France to O.M., and ANR ShootingStar (ANR-20-CE37-0018-04) to O.M. and B.C. T.B. was funded by a PhD fellowship from ENS.

## 6 Author contributions

S.E., T.B., O.M. and B.C. designed the study. T.B. and B.S.S. performed the experiments. S.E. and T.B. analyzed the data with help from O.M. and B.C.

S.E. did the model with help from T.B., O.M. and B.C. S.E., T.B., O.M. and B.C. wrote the paper.

